# Comparative reconstruction of the predatory feeding structures of the polyphenic nematode *Pristionchus pacificus*

**DOI:** 10.1101/2021.10.15.464383

**Authors:** Clayton J. Harry, Sonia M. Messar, Erik J. Ragsdale

## Abstract

*Pristionchus pacificus* is a nematode model for the developmental genetics of morphological polyphenism, especially at the level of individual cells. Morphological polyphenism in this species includes an evolutionary novelty, moveable teeth, which have enabled predatory feeding in this species and others in its family (Diplogastridae). From transmission electron micrographs of serial thin sections through an adult hermaphrodite of *P. pacificus*, we three-dimensionally reconstructed all epithelial and myoepithelial cells and syncytia, corresponding to 74 nuclei, of its face, mouth, and pharynx. We found that the epithelia that produce the predatory morphology of *P. pacificus* are identical to *Caenorhabditis elegans* in the number of cell classes and nuclei. However, differences in cell form, spatial relationships, and nucleus position correlate with gross morphological differences from *C. elegans* and outgroups. Moreover, we identified fine-structural features, especially in the anteriormost pharyngeal muscles, that underlie the conspicuous, left-right asymmetry that characterizes the *P. pacificus* feeding apparatus. Our reconstruction provides an anatomical map for studying the genetics of polyphenism, feeding behavior, and the development of novel form in a satellite model to *C. elegans*.

## 1 Introduction

A fundamental feature of nematodes used as models for developmental genetics is their constancy of cell number and fates (Sulston & Horvitz, 1977). Combined with a toolkit for genetics analysis, especially forward genetics, this feature allows the discovery of molecular interactions in and among cells of precise homology. As a classic example, vulva induction in *Caenorhabditis elegans*, the manipulation of individual cells and the factors they express revealed, in exquisite detail, how three different signaling pathways together pattern the fates of six precursor cells to produce a functional organ (Sternberg & Horvitz, 1989; Hill & Sternberg, 1992; Eisenmann et al., 1998). Leveraging such techniques to other nematodes, specifically those in which homologous cells could be identified, has since made cell-by-cell analysis available to comparative developmental genetics, especially in the nematode *Pristionchus pacificus* (Eizinger & Sommer, 1997; Sommer, 2009). Among the attributes of *P. pacificus* that benefit a comparative approach is its morphology. Unlike *C. elegans*, *P. pacificus* has a developmental polyphenism (*i.e.*, discontinuous plasticity) in its adult form, specifically in an evolutionary novelty, moveable teeth that enable predation (Fürst von Lieven & Sudhaus, 2000). Because this nematode can be used to study the proximal mechanisms of developmental plasticity (Bento et al., 2010; Ragsdale et al., 2013b), this system offers a route to studying the relationship between plasticity and morphological novelty as molecular processes among stereotypic, individual cells. Still, our understanding of such processes can only be as detailed as our anatomical understanding of the plastic trait.

The novel morphology of *P. pacificus* and other species of its family (Diplogastridae) consists of moveable teeth and associated armature in the mouth (*i.e.*, stoma, or buccal cavity). The polyphenism in *P. pacificus* is in the number and form of the teeth, as well as in the walls of the mouth, and the alternative morphs are induced in response to certain environmental cues (Bose et al., 2012; Serobyan et al., 2013, 2014; Werner et al., 2017). In *P. pacificus*, where the dimorphism has been best studied, the “wide-mouthed” (eurystomatous) morph has two opposing teeth and is omnivorous and predatory, taking a diet of bacteria, fungi, and other nematodes; the other, “narrow-mouthed” (stenostomatous) morph, which has only a single tooth, is strictly microbivorous (Wilecki et al., 2015; Lightfoot et al., 2019). The teeth and other mouth armature describing the polyphenism are extracellular, produced by both epithelial cells and myoepithelial cells (i.e., muscle cells that also secrete pharyngeal cuticle) (Wright & Thomson, 1981). Due to the inherent constraints of the nematodes’ tube-like body and tapered head, the cells making the mouth are complex in shape, notably containing long processes to where their nuclei are located (Hall & Altun, 2008). Therefore, unlike cells that are readily identified by light microscopy, such as the vulva precursor cells, cell identities in the head, mouth, and pharynx must be informed by transmission electron microscopy (TEM).

One feature of diplogastrid mouthparts especially deserves explanation from the underlying cellular architecture: their asymmetry, which defines the mouth of all species with the polyphenism (Kanzaki & Giblin-Davis, 2015). This asymmetry is both conspicuous (*i.e.*, not fluctuating; Ludwig, 1932; Palmer, 1996) and fixed for all known species with this morphology (*i.e.*, diretctional asymmetry *sensu* Van Valen, 1962). Specifically, the subventral tooth is always on the right, whereas the mouth’s left is equipped with a ridge of other structures, which vary by taxon. Even in *Pristionchus*, in which the stenostomatous morph has no subventral tooth, right and left always differ. The left side, like in the eurystomatous morph, always bears a row of denticles, albeit of different complexity between morphs (Ragsdale et al., 2015). In contrast, the closest outgroups to the family, including *C. elegans*, are strictly symmetrical in their cuticular feeding structures, both bilaterally and triradially (Chitwood & Chitwood, 1936). Symmetry breaking has been proposed as an origin of novel forms (Palmer, 2004), and indeed the teeth (and polyphenism) of Diplogastridae may have arisen together with asymmetry in the mouth (Susoy et al., 2015). A fine-structural map of the cells that produce these structures is needed to understand the developmental basis of the asymmetry, especially how it is achieved in animals with stereotypic cell fates.

The molecular basis for the mouth dimorphism of *P. pacificus* has enjoyed increasingly detailed attention, notwithstanding the limits of our knowledge of the head anatomy. In sensory cells, which are at the nematodes’ interface with the environment, factors that partially influence the dimorphism presumably transduce environmental information to an ultimate switch mechanism (Moreno et al. 2019; Lenuzzi et al. 2021). The polyphenism switch itself includes a series of enzymes, which are expressed in the central nervous system, presumably as part of an intercellular signaling system (Ragsdale et al., 2013b; Ragsdale & Ivers, 2016; Sieriebriennikov et al., 2018). At the end of this series is a sulfotransferase, which is expressed in the anterior epidermis (*i.e.*, face and mouth) and the pharynx (Bui et al., 2018; Namdeo et al., 2018). Also expressed in the pharynx, which produces the teeth, are two nuclear receptors that regulate mouth phenotypes. One of these, NHR-40, is expressed in pharyngeal muscle, and mutations in this gene show a total conversion between morphs, suggesting this transcription factor to be part of the switch itself (Kieninger et al., 2016). The other receptor, NHR-1, is expressed in both the muscle and glands of the pharynx, and receptor mutants include morphological defects that imply disruption of processes downstream of the switch itself (Sieriebriennikov et al., 2020). Some of the cells involved in this switch mechanism have been proposed based on similarity with *C. elegans* and on limited, unpublished data. However, precise identification of these and other cells, as well as any connectivity among them, still awaits their full reconstruction.

To date, the only complete reconstruction of the nematode face, mouth, and pharynx has been that of *C. elegans* (Albertson & Thomson, 1976; Wright & Thomson, 1981; White, 1988; Hall & Altun, 2008). However, a previous, fine-structural study of a species of Diplogastridae proposed homologies of mouth the cuticle (Fig. 1) and suggested that most cells of *P. pacificus* mouth should have identifiable homologs in *C. elegans* (Baldwin et al., 1997). If so, a complete reconstruction in *P. pacificus* would give the precision needed to pinpoint the intercellular differences relevant to the structures unique to Diplogastridae. One hypothesis, which is based on TEM observations of *Acrostichus* (= *Aduncospiculum*) *halicti* in that study, is that the teeth of diplogastrids are lined by an additional muscle cell layer missing in *C. elegans*, despite strict conservation of the non-muscular epithelia of the mouth in the two species (Baldwin et al., 1997). However, how many cell classes (typically, homologous pairs, triplets, or sextuplets of cells) and cell nuclei make up most of the rest of the diplogastrid mouth and anterior pharynx (corpus) has not been described, although they may also inform differences in the novel structure.

**Fig. 1.**
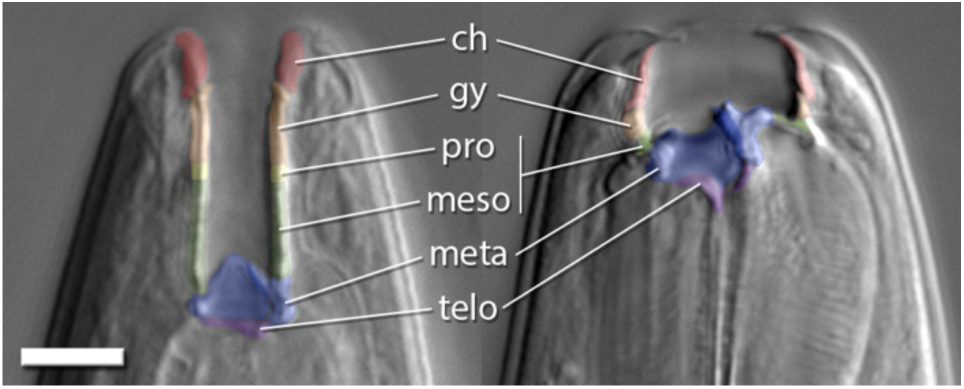
Proposed homologies of stoma (mouth) cuticle in *Caenorhabditis elegans* (“Rhabditidae”) and *Pristionchus pacificus* (Diplogastridae). Left image, *C. elegans*; right image, *P. pacificus*. False coloring indicates mouth regions: ch, cheilostom; gy, gymnostom; pro, prostegostom; meso, mesostegostom; meta, metastegostom; telo, telostegostom. Whereas cells producing cuticle in *C. elegans* are completely mapped, cell numbers and arrangements in *P. pacificus* were not previously described. Scale bar, for both images, 5 µm.

Partial TEM reconstructions of the head in nematodes outside of Rhabditina (*sensu* De Ley & Blaxter, 2002), the group that includes *C. elegans* and *P. pacificus*, enable comparative anatomy by polarizing changes between these two species. Homologies for the face epidermal (“hyp”) and interfacial (arcade) syncytia of *Acrobeles complexus* (Rhabditida: Tylenchina) and *C. elegans* are well supported: the number of syncytia and nuclei are identical between the two species, except for an additional, binucleate epidermal syncytium (“hypD”) in *A. complexus* (Bumbarger et al., 2006). The number of anterior pharynx cell classes is also similar among *C. elegans*, various Tylenchina species, and another putative outgroup, *Myolaimus* (Rhabditida: Myolaimina) (Van de Velde et al., 1994; Baldwin & Eddleman, 1995; De Ley et al., 1995; Giblin-Davis et al., 2010), although the homologies of some cells have been challenged based on cell lineage (Dolinski et al., 1998). Even the Tylenchina representative *Aphelenchus avenae*, despite having a protrusible, needle-like stomatostylet—another evolutionary novelty among nematodes—also shows high conservation of cells with *C. elegans*, supporting the reliability of outgroups for comparison. In *A. avenae*, the anterior pharynx is identical to *C. elegans* in the number of cell classes and nuclei, if not all their cell fates (Ragsdale et al., 2011). The number of facial and mouth syncytia are also identical, again with the single exception of hypD (Ragsdale et al., 2008, 2009; Ragsdale & Baldwin, 2010). These studies together suggest that, given a complete set of images through the same tissues in *P. pacificus*, cells of unequivocal homology can be determined for the face and feeding apparatus of this species.

A major step toward mapping the anatomy of *P. pacificus* was taken by Bumbarger et al. (2013), who reconstructed the connectome of its pharyngeal nervous system, a semi-autonomous “brain” within the nematode pharynx. That study found the pharyngeal connectome to comprise an identical number of neurons (20) in *P. pacificus* and *C. elegans*, although its wiring has diverged in ways that reflect behavioral differences between the species. Importantly for comparative nematode anatomy, this work yielded a practically complete set of TEM images through the entire head and neck of *P. pacificus*. This image set has since been used to reconstruct the three pharyngeal gland cells of *P. pacificus* (Riebesell & Sommer, 2017) and, outside of the pharynx, the anterior sensory cells and organs of the species (Hong et al., 2019). Here, we used the images to three-dimensionally reconstruct all 74 epithelial and myoepothelial cells of the face, mouth, and pharynx. By completing the anatomical map of the *P. pacificus* feeding apparatus, we identified fine-structural features—including in cell arrangement, connectivity, and symmetry—that underlie the novel feeding morphology of *P. pacificus*.

## 2 Materials and Methods

Reconstructions were performed from a set of 2662 serial, 50 nm-thick TEM sections through the head of an adult, eurystomatous *P. pacificus* hermaphrodite (*i.e.*, self-fertilizing, morphological female) of *P. pacificus* that was high-pressure frozen and then fixed by freeze-substitution. Preparation for TEM, imaging of sections, and ordering and alignment of images were performed previously, as described by Bumbarger et al. (2013). The image series (“Series 14”) used was of Specimen 107 from that study, the specimen since used to reconstruct the pharyngeal gland cells (Riebesell & Sommer, 2017). Because the number of anterior epithelial and muscle cells is invariant within species in all Rhabditida (Rhabditina, Tylenchina, and Myolaimina) sufficiently examined, and because in most cases these numbers are conserved even across species, we performed our reconstruction from one individual. Throughout the image series, individual contours were manually traced for volume segmentation using TrakEM2 (Cardona et al., 2012). For each cell, all contours in all unobstructed images in the alignment were annotated. Volume segmentation was performed for the contours of the epidermal syncytia identified as hyp1 through hyp4, the arcade syncytia, all cells and syncytia in the pharynx except neurons and glands, the extracellular cuticle of the face and teeth, and an atypical cell hypothesized to one of the two XXX cells. Volumes for nuclei of all cells and syncytia were additionally segmented. Fully segmented volumes were transferred to Blender v. 2.93 (blender3d.org), in which automatic meshing errors were manually corrected according to the original TEM data and volumes were visualized. Individual object files for the volumes, which can be manipulated for visualization using various software platforms, have been deposited in Dryad (datadryad.org).

To present TEM data, original micrographs were automatically stitched by globally optimized registration (Preibisch et al., 2009), as implemented in ImageJ. For fine-structural observations of *C. elegans*, original TEM images were accessed through WormImage (wormimage.org).

Reconstructed anatomy of *P. pacificus* follows conventional names in *C. elegans*, as summarized by Hall & Altun (2008). Comparison of the *P. pacificus* pharynx to other nematodes is based on previously proposed homologies as summarized and assigned *C. elegans* nomenclature by Ragsdale et al. (2011).

## 3 Results

Our reconstruction of the anterior epithelia of *P. pacificus* revealed that every cell, syncytium, and nucleus of the facial epidermis and feeding apparatus could be reliably named based on homologs in *C. elegans* (Fig. 2). Inferences of homology are based on relative position, numbers of nuclei per cell class, and cell fate (*i.e.*, epithelial *vs*. myoepithelial), all of which were identical with those of *C. elegans*.

**Fig. 2.**
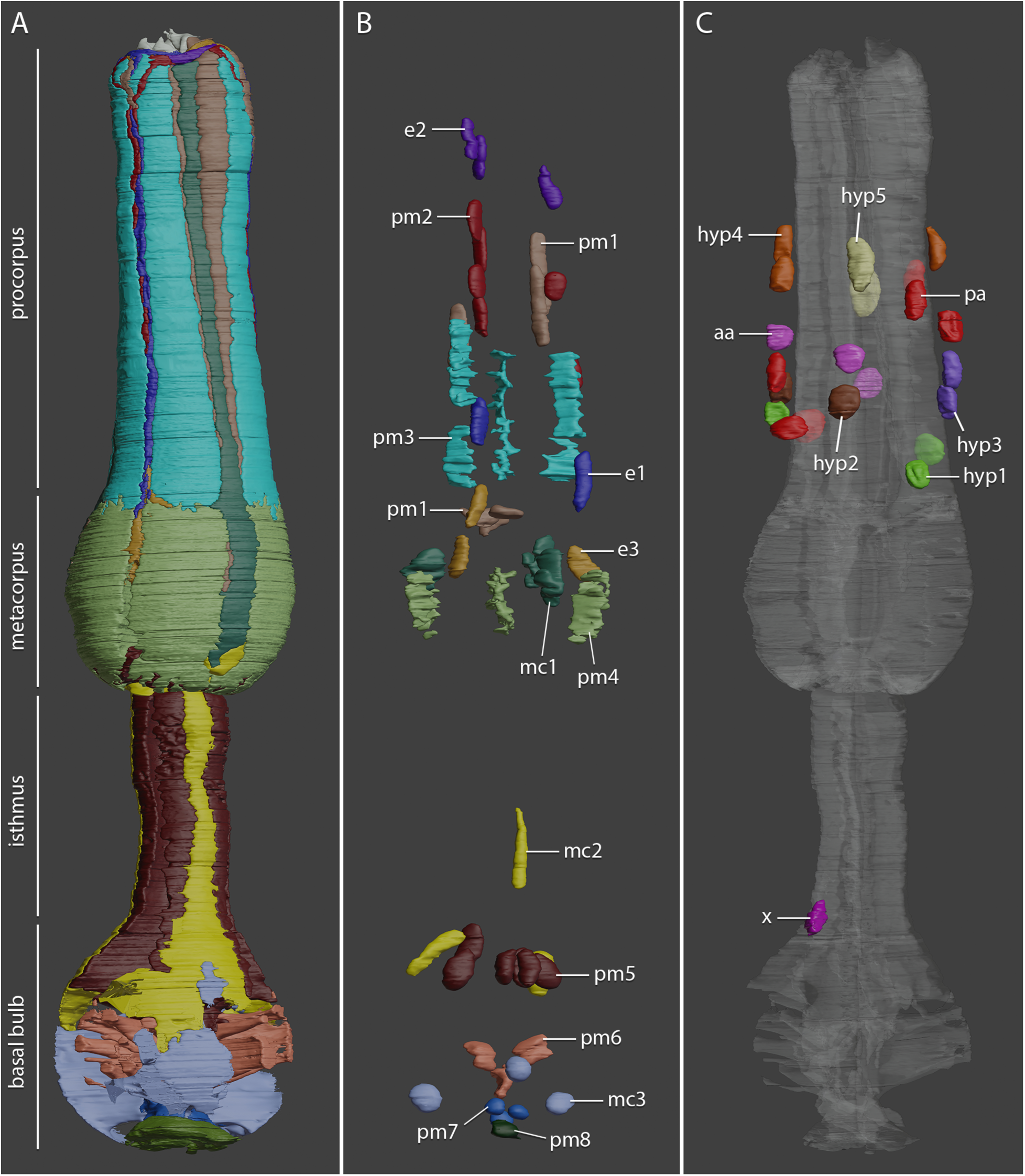
Three-dimensional models of complete pharynx epithelia, their nuclei, and epidermal nuclei of *Pristionchus pacificus*. Cell classes distinguished by color. A, Pharyngeal epithelial cells and syncytia, in left lateral view. Regions labeled define the corpus (procorpus + metacorpus) and postcorpus (isthmus + basal bulb). Colors of cell/syncytial membranes follow (B). B, Nuclei of pharyngeal epithelia, in left lateral view. Labels shown for one nucleus per class, except pm1, labeled for both procorpus-type and metacorpus-type nucleus positions. C, Anterior epidermal and arcade syncytial nuclei, in left lateral view, with pharyngeal muscle shown (transparent white) for spatial context. Labels shown for one nucleus per class. Abbreviations: aa, anterior arcade; pa, posterior arcade; x, standalone cell hypothesized to be XXXL.

### 3.1 Face and anterior mouth epithelia

The face of *P. pacificus*, as in *C. elegans* and more distantly related microbivores, is lined by a series of ring-like syncytia with posteriorly directed processes (“somal extensions”) ending in cell bodies containing nuclei (Figs 2C, 3A). These extensions are bundled together in four groups of extensions: dorsal, left and right dorsolateral, and ventral. Within each bundle, every somal extension was confirmed to make direct contact, along some part of its length, with every other extension in the same bundle. The nature of this connectivity is not often clear, although we hypothesize that these and other observed intercellular contacts include gap junctions— specifically, Type II gap junctions as defined for *P. pacificus* (Bumbarger et al., 2013)—which we could not always objectively detect at the resolution offered by the data.

**Fig. 3.**
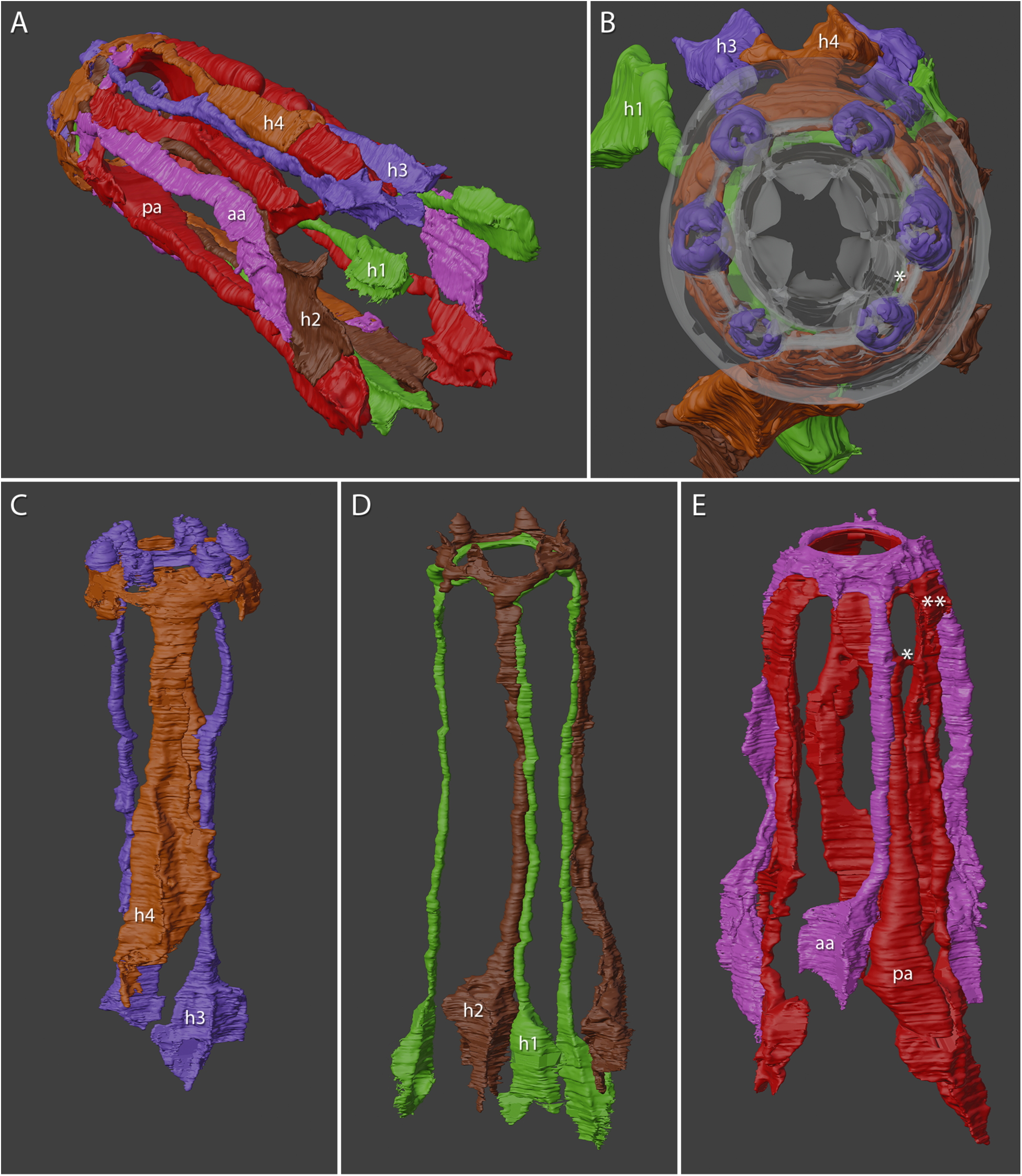
Three-dimensional models of face and mouth epithelia of *Pristionchus pacificus*. Colors follow Fig. 2C. A, Membranesreconstructed arcade and anterior epidermal syncytia, shown in oblique, left posterior view. B, Epidermal syncytia hyp1–hyp4 and anterior cuticle (white, transparent), viewed *en face*, with dorsal above. hyp4 crosses over lip syncytia hyp2 and hyp3 medially to line mouth flaps. Star, left ventrolateral hyp4 extension. (C)–(E) in identical orientation, in slightly oblique ventral view. C, hyp3 and hyp4. D, hyp1 and hyp2. E, Anterior and posterior arcade syncytia. Star, bridge between left ventral somal extension (not connected to toroid) and ventral somal extension. Double star, left subventral pseudosomal extension.

In *P. pacificus*, the syncytial rings of hyp1 through hyp4 and the arcade syncytia are thin and telescoped together such that their syncytial rings have parts that adjoin every other of the six syncytia (Figs 3B, 4A, B). Although they generally line different transverse sections of the mouth cuticle, the margins that some line are very thin. The facial syncytium most distal from the mouth opening is hyp4, which lines the anterior body wall cuticle (Fig. 4A). The body wall cuticle posterior to this syncytium is lined by hyp5, which forms a sheet-like collar around the head. Although the cell membrane of hyp5 is not reconstructed here, we report the positions of its two nuclei, as these are relevant to the map of anterior epithelia. Like in *C. elegans*, the hyp5 nuclei are lateral and at about midway along the pharyngeal procorpus (Fig. 2C).

**Fig. 4.**
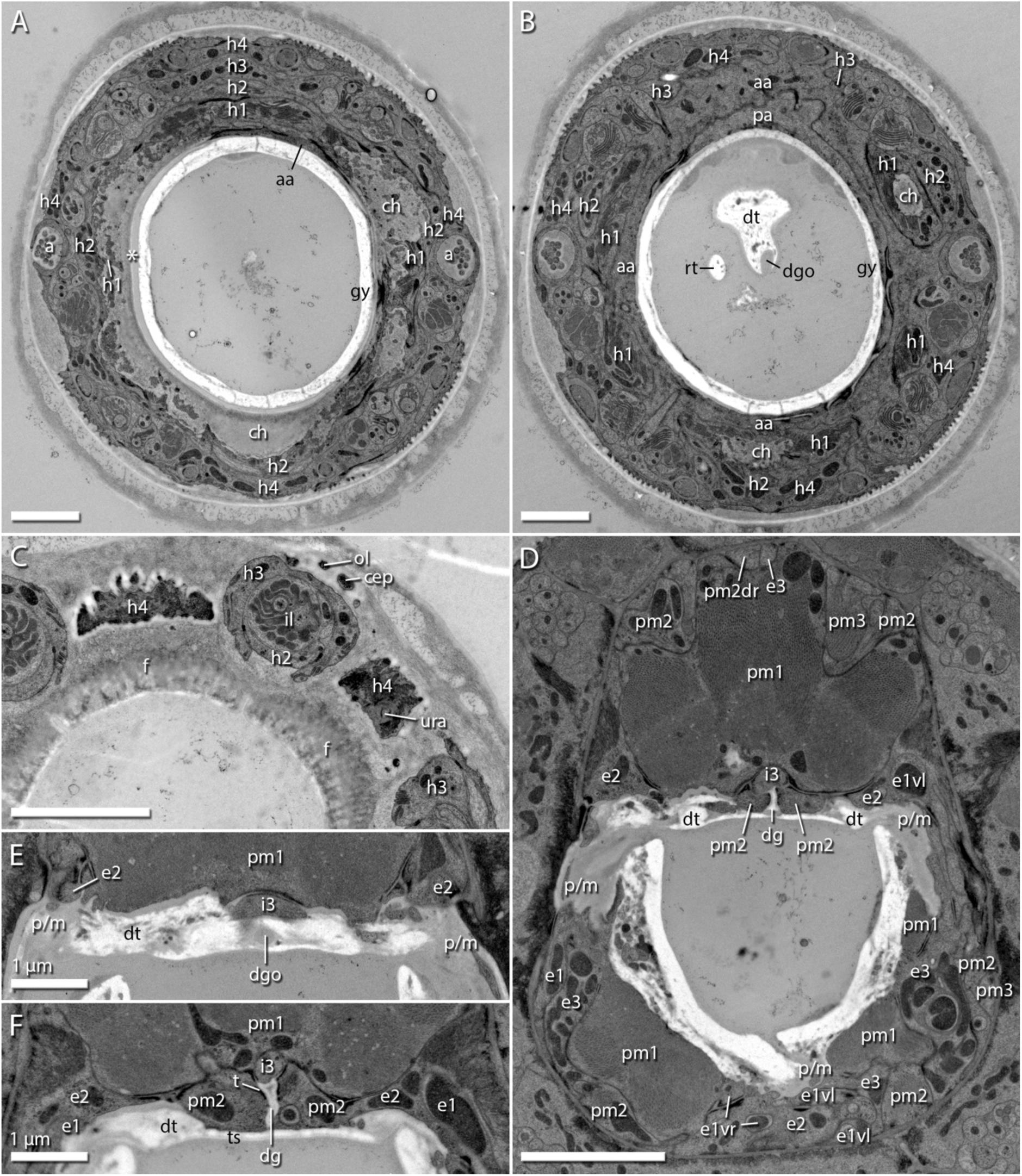
Transmission electron micrographs of transverse sections through the face and mouth of *Pristionchus pacificus*. Top of each image is dorsal. Scale bars 3 µm unless otherwise indicated. A, Section through cheilostom (ch) and gymnostom (gy). hyp1 (h1) and hyp2 (h2) secrete cheilostom. hyp4 (h4) lines anterior body wall, including at amphid (a) openings, and is associated with sensory neuron URA (ura). hyp1–hyp4 are telescoped together as shown dorsally. Star, external space. B, Section through gymnostom and tips of dorsal tooth (dt) and right subventral tooth (rt). Section shows both anterior arcade (aa) and posterior arcade (pa) syncytia lining gymnostom; somal extensions of hyp1–4 also shown. C, Section through labial and anterior mouth. hyp2 and hyp3 form lips and are associated with several sensilla, *e.g.*, cephalic sensillum neuron CEP (cep), inner labial sensillum (il); outer labial sensillum neuron OLQ (ol). hyp4 traverses between lips to line flaps (f) of mouth opening. D, Section through right subventral tooth, left subventral ridge of denticles (lr), and cuticle posterior to dorsal tooth. Dorsal pm2 muscles insert on dorsal gland (dg) where it meets mouth cuticle. Opposite dorsal tooth, ventral pm1 muscles insert on subventral tooth and ridge. Subdorsal and ventral apices of mouth comprise electron-lucent cuticle of pro- and mesostegostom (p/m), lined by e1, e3, and e2; complexity of their arrangement shown here by forms of e1VR (e1vr), e1VL (e1vl), and ventral e2 cells. E, Section through base of dorsal tooth (dt), anterior to section in (D). Dorsal gland orifice (dgo) runs through tooth. F, Section posterior to base of tooth, just posterior to section in (D). Dorsal pm2 cells, together with pharyngeal interneuron I3 (i3), adjoin dorsal gland cuticle, to which tonofilaments (t) of pm2 cells attach.

In addition to producing the anterior body wall cuticle, hyp4 also lines the flaps of the mouth, structures unique to *Pristionchus* and other diplogastrid genera. Specifically, the hyp4 ring has medially directed extensions that cross over the adjoining hyp3 syncytium anteriorly (Figs 3B, 4C). Thus, the flaps of the mouth are homologous with the body wall cuticle rather than lip or mouth cuticle. The four sublateral of these extensions enclose the termini of the URA sensory neurons, which were identified previously (Hong et al., 2019). The hyp4 and hyp3 syncytia together surround the amphids, cephalic sensilla, and outer labial sensilla where they meet the body wall. Posteriorly, hyp4 has two somal extensions, one dorsal and one ventral (Fig. 3C): the former ends in a cell body containing two, longitudinally arranged nuclei, while the latter contains one nucleus. Together, the hyp4 nuclei are the most anterior nuclei of all epithelial cells, or cells of any kind outside of the nematode pharynx. The remaining four of the six radial sectors of the hyp4 ring show short processes that contact the posterior arcade syncytium, while the stretches between those processes contact the anterior arcade syncytium.

Syncytium hyp3 produces the anterior and distal (outer) cuticle of the lips, as well as the anterior most rim of the cheilostom, the anterior most capsule of the mouth itself (Figs 3B, 4C). The syncytial ring thinly connects six bulges within each of the lips, of which there are two subdorsal, left and right lateral, two subventral. Each of these bulges is grooved medially and has a hollow center, through which the inner labial sensilla pass (Figs 3C, 4C). Excluding URA, the anterior, internal sensory neurons, which were described previously (Hong et al., 2019), also terminate in pockets of hyp3. This syncytium has two subdorsal somal extensions (Fig. 4B), the nuclei of which are posterior to and on either side of the dorsal nucleus of the posterior arcade cell. These somal extensions run for some distance apart from those of other syncytia, although the hyp3 extensions connect with the other dorsal extensions (hyp4, anterior arcade) posteriorly along their length (Fig. 3A).

The hyp2 syncytium lines a ring of mouth cuticle just posterior to hyp3. Further, it extends six processes anteriad to produce the medial (inner) cuticle of the lips (Fig. 4C). Thus, hyp2 and hyp3 together bound the anterior sensilla of the face. As in other Rhabditida comparably described (Bumbarger et al., 2006; Hall & Altun, 2008; Ragsdale et al., 2008), the somal extension of hyp2 are asymmetrical in that there is one left dorsolateral and one ventral somal nucleus; only a short posterior process (“pseudosomal extension”) is present dorsolaterally on the right (Fig. 3D). The cell bodies of hyp2 posteriorly adjoin ventral and left dorsolateral cell bodies of the anterior arcade syncytium.

The posterior margin of the cheilostom is lined by hyp1, which has the thinnest of the syncytial rings and, together with hyp2, the thinnest somal extensions (Fig. 3D). Each of the three somal extensions, which are dorsolateral and ventral, run inside of a shallow groove of an extension of the posterior arcade syncytium; the ventral and left dorsolateral extensions additionally run together with those of hyp2 (Fig. 3A). As in *C. elegans*, the two dorsolateral hyp2 cell bodies are more posterior than the ventral, which lies next to the ventral hyp2 nucleus, and are the most posterior of all syncytia described herein. Of the cell bodies for hyp1– hyp4 and the arcade syncytia, the two subdorsal ones of hyp1 are unique in that they do not contact any cell bodies of other epithelia.

### 3.2 Internal mouth capsule and arcade syncytia

In *Pristionchus* and several other diplogastrid genera, the mouth contains a tube, homologous with the gymnostom of other nematodes, that is concentric within the outer wall of the mouth. The posterior base of this tube in *P. pacificus* is lined, for a thin margin at its base, by the two thin epithelial rings of the arcade syncytia (Figs 3E, 4B). The anterior arcade syncytium, which adjoins hyp1 anteriorly, has three robust somal extensions—two dorsolateral and one ventral— each with one nucleus. The posterior arcade syncytium, which adjoins the pharynx posteriorly, has six nuclei. The ventral and ventrolateral of these are more posterior than the dorsal and dorsolateral (Fig. 2C, 3E). The somal extensions of the posterior arcade syncytium are each associated with one of the four bundles of somal extensions, except for the left and right ventrolateral processes, which run unaccompanied to cell bodies that then contact cell bodies of other syncytia (Fig. 3A).

Although there are six posterior arcade nuclei distributed around the circumference of the nematode, the processes leading to their cell bodies are radially asymmetrical. Specifically, some processes end as pseudosomal extensions (*i.e.*, without nuclei), others split posteriorly to contain two cell bodies, and still others are bridged together. Similar asymmetries of the posterior arcade syncytium of *C. elegans* apparently differ between specimens, given the description by Hall & Altun (2008) compared with that by Wright & Thomson (1981). Therefore, we presume differences likewise occur within *P. pacificus*. Nevertheless, we report the particular pattern observed for Specimen 107. First, only the right ventrolateral and left dorsolateral somal extensions are simple, without branches or bridges. The left ventrolateral process is a pseudosomal extension, while the ventral process splits just posterior to the latter’s end into two long extensions to the ventral and left ventrolateral nucleus (Fig. 3E), the cell bodies for which adjoin but are not syncytial where they meet. The dorsal and right dorsolateral extensions form a thin bridge about halfway along their length (Fig. 3E).

### 3.3 Teeth and pharyngeal corpus

The region of the pharynx lining the mouth (stegostom) comprises several classes of cells that are complex in their relative positions and in where they line the mouth cuticle, including the teeth (Fig. 2A). These cells all line a relatively short length of the anterior pharynx, such that all of their nuclei are located at the end of processes extending posteriad into the pharynx (Figs 2A, 3B).

The most anterior cell classes of the pharynx are the “e” (epithelial) cells: e1 and e3 line thin strips of cuticle that attach to the two teeth and left subventral ridge to the gymnostom (Fig. 5A). Together, this cuticle blends the prostegostom and mesostegostom. These strips of cuticle also extend posteriad into the radii, or apices, of the pharyngeal lumen, that separate each of those three structures. Both e1 and e3 cells line electron-lucent cuticle distinct from that of the tooth and more posterior regions of the mouth (Fig. 4D). The e1 and e3 are classes of three radial cells each: one dorsal and two (left and right) subventral, which are interspaced along the cuticle by the e2 cells (Fig. 6). Each e1 and e3 cell extends a process posteriad through the anterior pharynx (procorpus) muscle (pm3). The cell bodies of the e1 cells span the procorpus to the median bulb (metacorpus), whereby the e1VR and e1VL nuclei are in the procorpus and the e1D nucleus is more posterior, at the transition to the metacorpus (Fig. 2B).

**Fig. 5.**
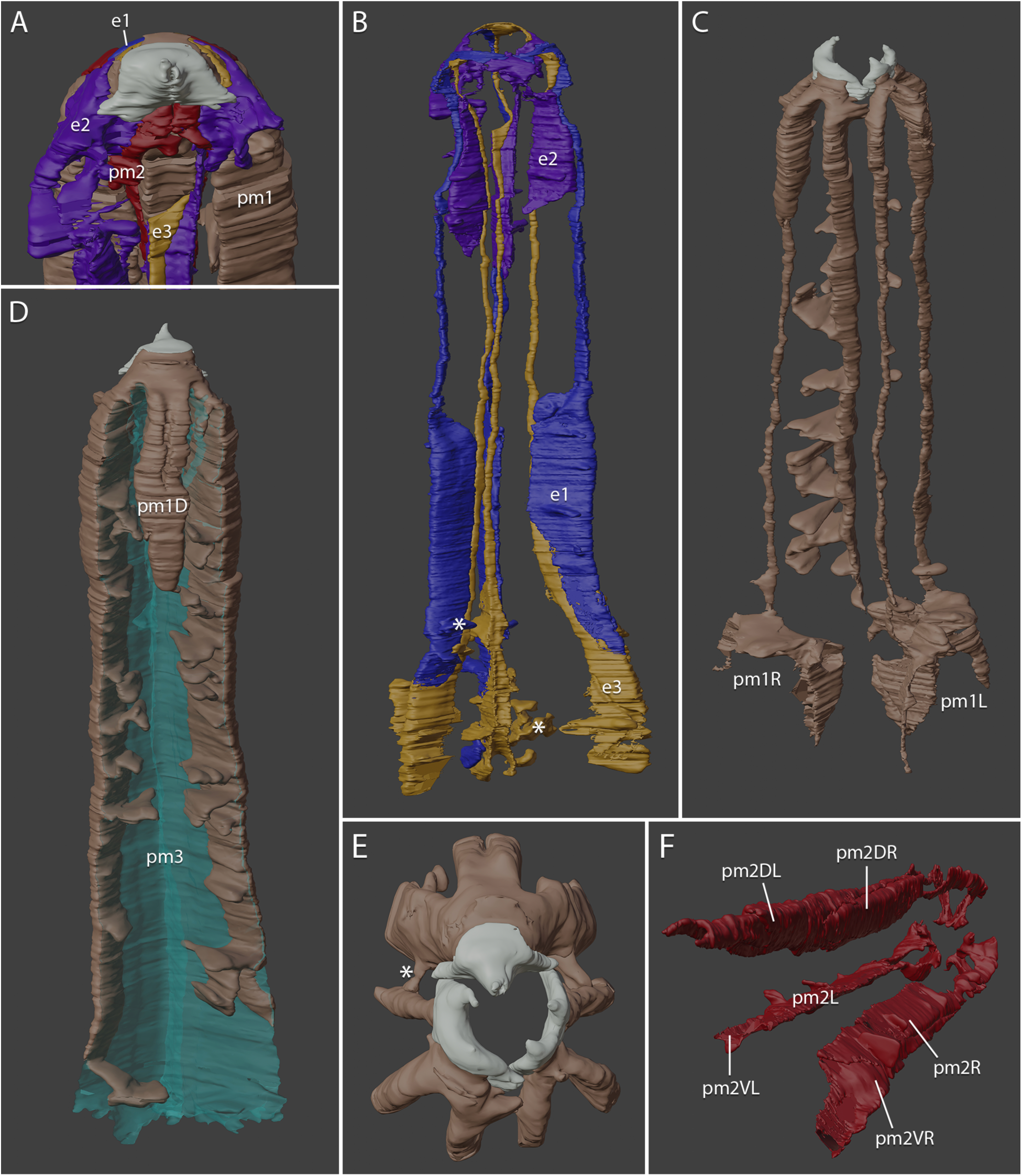
Three-dimensional models of pharyngeal mouth armature and associated muscle and epithelial cells of *Pristionchus pacificus*. Colors follow Fig. 2B. A, Dorsal tooth (white), dorsal pm1 muscle, dorsal pm2 muscles, and epithelial cells e1D, e2DL, e2DR, and e3D, shown in oblique ventral view. Cells e1–e3 wrap around tooth, which is lined dorsally by pm1 muscle. Dorsal pm2 muscle cells extend anteriad within base of tooth. B, cells of e1, e2, and e3, in slightly oblique ventral view. Stars indicate examples of transverse processes of e1 and e3. C, Right subventral tooth (white), left subventral ridge (white), and associated pm1VR and pm1VL muscles, respectively, in slightly oblique ventral view. Ventral arm of pm1VR has many transverse processes and terminates in procorpus, unlike its medial arm and both arms of pm1VL, which are thinner, simpler, and terminate in metacorpus. D, Dorsal tooth, pm1D, and pm3D muscle (transparent), in slightly oblique dorsal view. Both adradial arms of pm1D are large, extend transverse processes into pm3D, and terminate in procorpus. Short radial arm of pm1D is also larger than in other pm1 cells. E, Both teeth, left subventral ridge, and anterior parts of attached pm1 muscles, viewed *en face*, with dorsal above. Processes connect a large adradial muscle arm of one pm1 syncytium to small arm of neighboring pm1 syncytium. Star, left subdorsal process of pm1D. F, pm2 cells, in oblique, right posterior view.

**Fig. 6.**
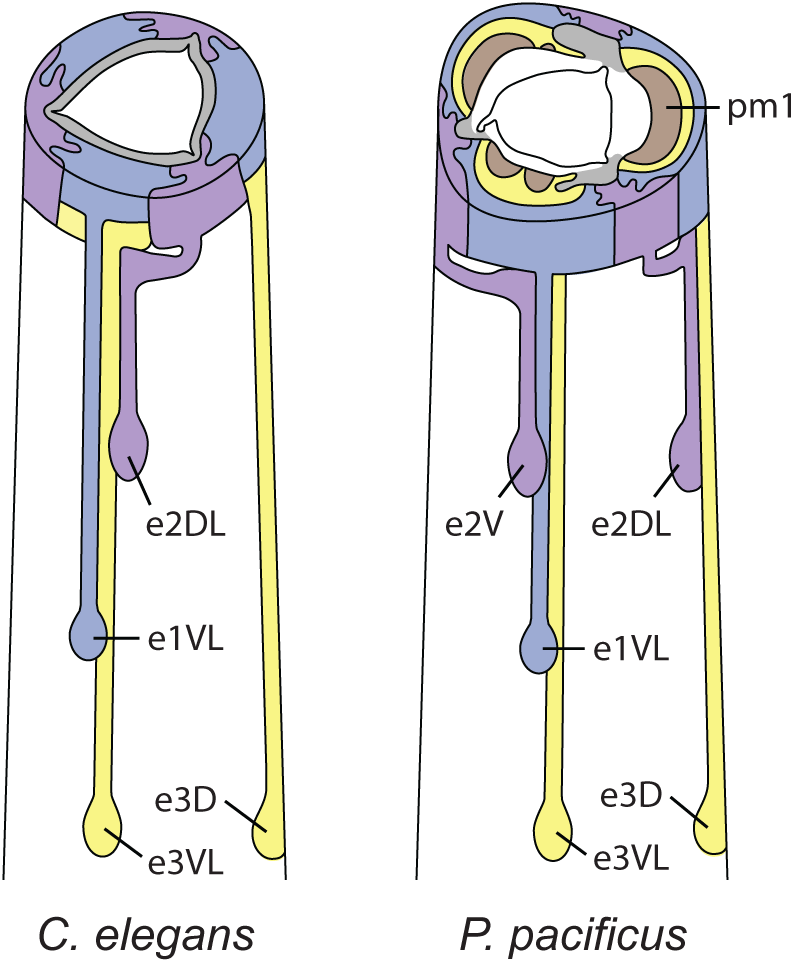
Stylized diagram of cells e1, e2, and e3 in *Caenorhabditis elegans* and *Pristionchus pacificus*. Depiction of cells in *P. pacificus* is based on the individual described herein (Specimen 107). Grey indicates prostegostom (*C. elegans*) and pro- and mesostegostom (*P. pacificus*). Arrangement of e cells is complex in *P. pacificus*, such that they overlap with tooth and other metastegostomatal structures lined by pm1 (brown). All three e cells of *P. pacificus* additionally line cuticle anterior to that shown in diagram. Diagram of *C. elegans* modified from Albertson & Thomson (1976). Relative cell sizes not to scale.

The e3 cells have their cell bodies and nuclei entirely within the metacorpus (Fig. 2B). Each e3 cell body has a large area of contact with the e1 cell body, and in the case of the dorsal e1 and e3 cells, the two cell bodies interdigitate substantially (Fig. 5B). The cell bodies of all e1 and e3 cells have several transverse processes (Fig. 5B), which extend into the adjoining pm3 muscle cells.

The two (left and right) subdorsal and ventral apices of the mouth are lined by three e2 cells, which also line the pharynx more posteriorly than e1 and e3 (Figs 4D, E, F 5A, B), specifically the cuticle between the muscles inserting on and just posterior to the teeth (pm1, pm2). The e2 cells are relatively short, such that they contain the most anterior nuclei of the pharynx (Figs 2B, 5B). As in *C. elegans*, the e2 nuclei are not located in the same sector as where they line the pharynx, although the pattern is mirror-image to that in *C. elegans*: in the observed specimen of *P. pacificus*, the e2DL nucleus is dorsal, the e2DR nucleus is right subventral, and the e2V nucleus is left subventral (Fig. 6). Of these, the dorsal nucleus (e2DL) is posterior to the other two. Like the e1 and e3 cells, the e2 cells send transverse processes into the pm1 cells, and these processes are more numerous on e2DL and e2DR than in e2V (Fig. 5B).

Posterior to the e1 and e3 cells are the pm1 muscles, which comprise three binucleate syncytia, one dorsal and two subventral (Figs 2A, B, 5C, D). The pm1 muscles insert on the dorsal tooth, right subventral tooth, and left subventral ridge of denticles (Figs 4D, E, 5E). All three cells consist of two long adradial arms, each containing a nucleus posteriorly, between which there is a shorter, central (radial) arm (Figs 4D, E, 7A). Whereas the adradial arms are likely homologous with the nucleus-containing processes of pm1 in *C. elegans* and microbivorous Tylenchina, the radial arms are unique to *P. pacificus* with respect to those taxa (Baldwin et al., 1995; De Ley et al., 1995). All pm1 arms contain longitudinal myofilaments (Figs 4D, F, 7A), which extend posteriad to about where the shorter processes end. Of the pm1 muscles, pm1D, which inserts on the dorsal tooth, is by far the largest in its class. It is this syncytium that explains the dorsoventral anisomorphy of the pharynx, a feature easily seen by light microscopy. The adradial arms of pm1D are both thick in cross section and along their lengths send numerous large, transverse extensions into the adjacent pm3 muscle (Figs 5D, 7B). Further, there is a difference in the form and size between the two subventral pm1 cells that articulates the bilateral symmetry of the gross morphology. Specifically, pm1VR—the muscle that inserts on the moveable, right subventral tooth—has an adradial arm (subventral) similar in form to those of pm1D, as well as an adradial arm (lateral) that is thin and lacks those processes. In other words, the large pm1VR arm is in opposition to the pm1D cell, presumably actuating, in large part, the right subventral tooth. In contrast, pm1VL, which inserts on the immoveable, left subventral ridge only has two thin arms. Further, this asymmetry correlates with a distinct dimorphism in nuclei: those in the large arms (one in pm1D, two in pm1VR) are all long and located in the procorpus, and their cell bodies terminate in the procorpus, whereas nuclei in the thin arms are short and end in cell bodies that are completely within the metacorpus (Figs 2B, 5C, D, 7B). Related to this asymmetry is how the three pm1 cells all make contact with one other, which they do through small transverse processes that run between e2 and the anterior margin of mc1 (corpus marginal cell), the cells that otherwise separate the pm1 cells (Figs 2A, 5E, 7A). These processes each extend from a large pm1 adradial arm (two in pm1D, one in pm2) to the thin adradial arm of the adjacent syncytium.

**Fig. 7.**
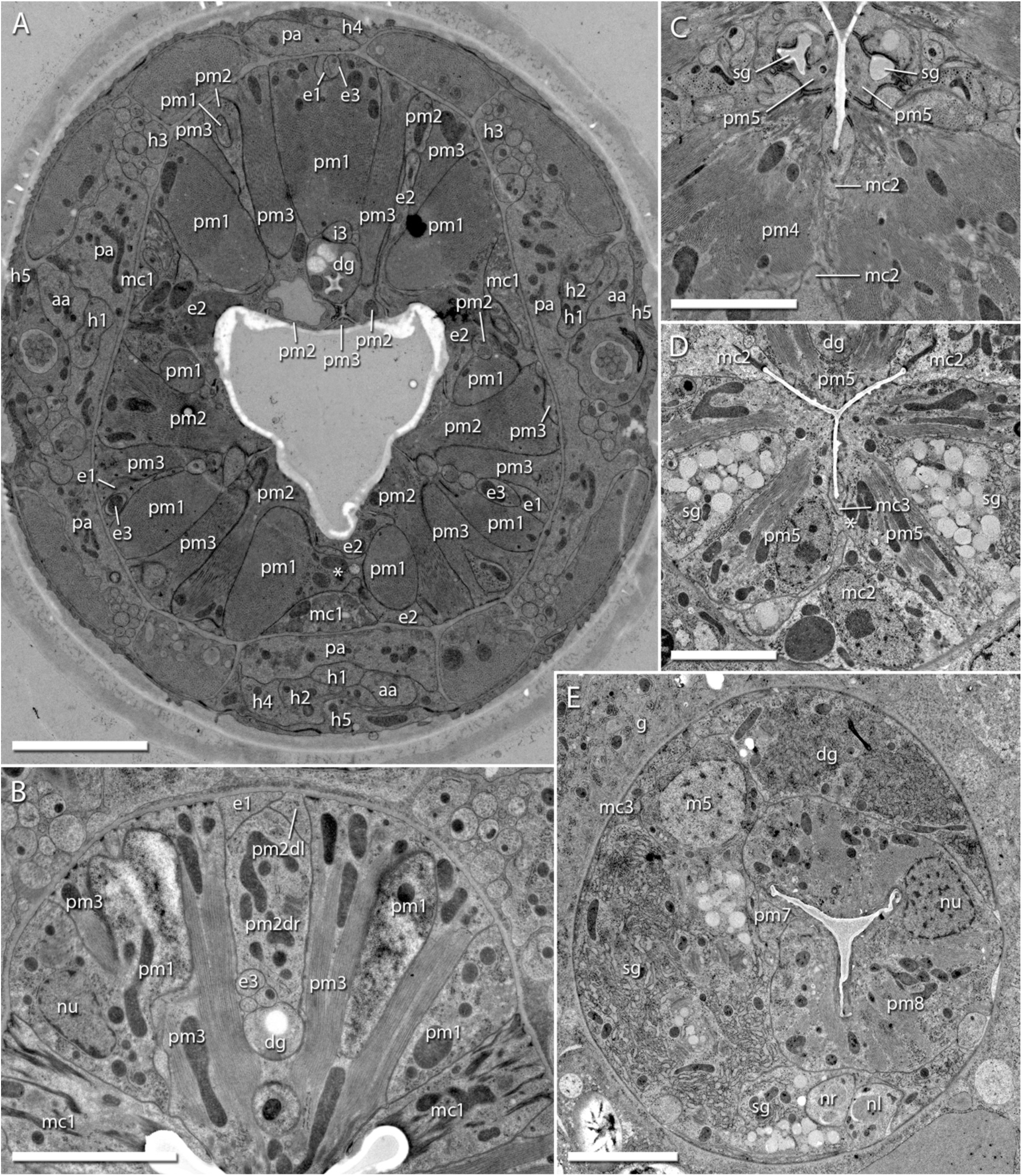
Transmission electron micrographs of transverse sections through the pharynx of *Pristionchus pacificus*. Top of images is dorsal. Images ordered from anterior to posterior. Scale bars 3 µm. A, section through telostegostom and surrounding tissues. Each pm1 syncytium branches into three arms, one radial and two adradial. Ventral arms of pm1VR and pm1VL connect through a transverse process of former (star). Anterior reaches of mc1 and dorsal pm3 cells also shown. Outside pharynx, somal extensions of arcade and anterior epidermal syncytia run in bundles; lateral parts of posterior arcade (pa) syncytium are shown anterior to where they split into somal extensions. Sheet-like hyp5 (h5) syncytium labeled at its ventral and two lateral swellings. B, Section through middle of procorpus, showing dorsomedial transverse processes of pm1 enclosed by pm3. Large adradial arms of pm1 also each have a nucleus (nu) in procorpus. C, Section though posterior metacorpus, near where subventral glands (sg) empty into pharyngeal lumen. A gap separates mc2 where it lines lumen from where its attaches to basal lamina of pharynx, accommodating a bridge between subventral sectors of pm4. D, Section through posterior part of isthmus. A gap separates mc3, lining lumen, from mc2, along basal lamina; subventral sectors of pm5 are syncytial through this gap. E, Posterior end of basal bulb, showing cap-like pm8 cell with its single nucleus (nu), posterior end of right subventral pm7 cell and right subdorsal mc3 cell. Gut (g) and pharyngeal neurons M5 (m5), NSML (nl), and NSMR (nr) labeled for spatial context. Abbreviations follow Fig. 4.

The pm2 cells consist of three pairs of adradial, uninucleate cells (Fig. 5F, 7A). The muscles have small insertions on the walls of the mouth (telostegostom) just posterior to the teeth and denticle ridge (Figs 4D, F, 5A). Where the dorsal pm2 cells insert on the cuticle, they abut either side of the entry of the dorsal gland into the cuticle at the base of the tooth (Fig. 4D), and their myofilaments run distally from the dorsal gland duct, to which the dorsal pm2 cells are secured through tonofilaments (Fig. 4F). The cell processes of all pm2 cells run along the short, radial processes of the pm1 cells and, posterior to that, to each other. Like the pm1 muscles, the pm2 cells have transverse extensions into pm3, although these are far fewer than for the large adradial arms of the pm1 cells. The cell bodies of pm2 are within the procorpus, lying in distal grooves in pm3 (Fig. 2A), with each pair of cell bodies in longitudinal series (Fig. 5F). Although the cell bodies in each pair adjoin each other, they are not syncytial. The dorsal pair of cell bodies is posterior to the subventral pairs (Fig. 2B).

The pm3 and pm4 cells make up the pumping (dilating) muscles of nearly all of the procorpus and metacorpus, respectively (Fig. 2A). The pm3 nuclei are located in the posterior half of their cells, and the pm4 nuclei are near the circumference of the metacorpus at its widest (Fig. 2B). The pm3 muscles comprise three radial syncytia of two adradial nuclei each and, as described above, accommodate processes from e1, e2, e3, pm1, and pm2. As described by Burr & Baldwin (2016) for *C. elegans*, pm3 in *P. pacificus* might more accurately be considered “paired cells”—that is, cells retaining their own cytoskeleton but connected by a common cytoplasm—than syncytia, although we retain the latter term here for consistency with other nematode reconstructions. The pm4 muscle, which is mostly divided into three sectors around the pharynx, is a single syncytium with three pairs of adradial nuclei (Fig. 2B).

The three sectors of pm4 connect through relatively small bridges, two subdorsal and one ventral (Figs 7C, 8B). Between pm4 and the next posterior set of muscles (pm5) is the pharyngeal nerve ring of the pharynx, which is described elsewhere (Bumbarger et al., 2013; Bumbarger & Riebesell, 2015).

**Fig. 8.**
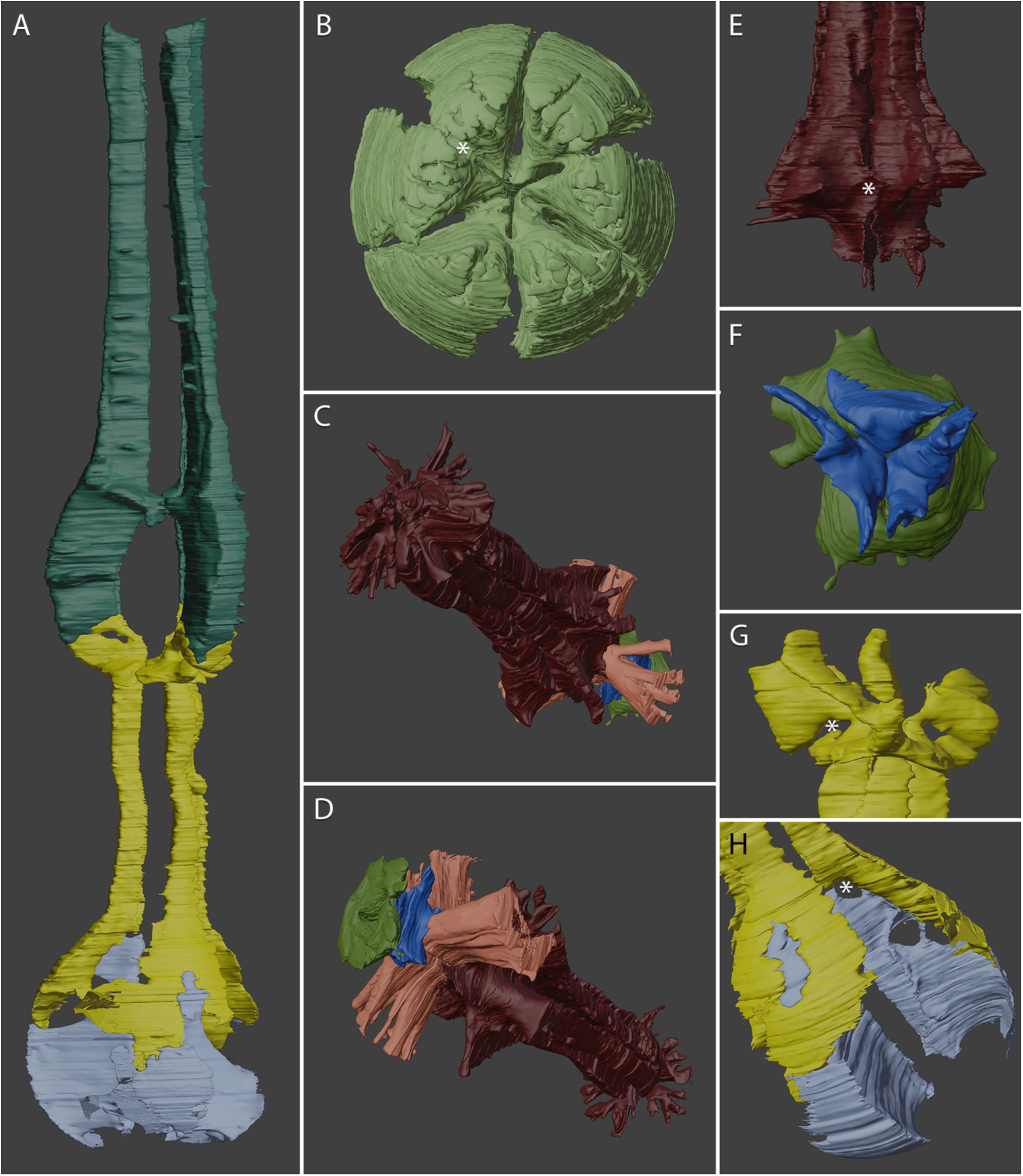
Three-dimensional models of marginal cells, metacorpus, and postcorpus of *Pristionchus pacificus*. Colors follow Fig. 2B. A, Marginal cells mc1, mc2, and mc3, in left lateral view. Both mc1 and mc2 completely surround lumen in metacorpus. B, Syncytium pm4, in posterior view, with dorsal above. Bridges join dorsal and two subventral sectors into a syncytium; star, left subdorsal bridge. C, Postcorpus muscles pm5, pm6, pm7, and pm8, in oblique, left anterior view. D. Muscles pm5–pm8 in oblique, right posterior view. E, pm7 and pm8 muscles in anterior view, with dorsal above. F, Posterior part of pm5 syncytium, in ventral view. Star, bridge between subventral sectors of pm5. G, Anterior part of mc3 marginal cells, in dorsal view. Star, groove in mc2DL through which pm4 forms syncytial bridge. H, Marginal cells mc2 and mc3, in oblique, left ventral view. Star, groove in m32DL through witch pm5 forms syncytial bridge.

Lining the radii of the procorpus and most of the metacorpus are the three, non-muscular mc1 marginal cells (Figs 2A, 7B, 8A). These cells, each with a single nucleus in the metacorpus, line the radii (*i.e.*, the subdorsal and ventral folds of the relaxed lumen) of the pharynx, each one as an anchor to a muscle cell opposing it along the pharyngeal lumen. Where the two subventral glands enter the pharyngeal lumen in the metacorpus, the mc1 cells completely surround the lumen in a ring, thereby spacing the pm3 cells anteriorly from the pm4 cell posteriorly (Fig. 8A).

### 3.4 Pharyngeal postcorpus

The postcorpus, the peristaltic part of the pharynx, is made up of a longitudinal series of muscle cells anchored by marginal cells (Figs 2A, 8C, D) and, in the basal (terminal) bulb, three large gland cells that are described elsewhere (Riebesell & Sommer, 2017). Distinct from in *C. elegans*, but similar to the outgroup *Aphelenchus avenae* (Ragsdale et al., 2011), the anterior part of the pharyngeal isthmus and surrounding basal lamina is telescoped within the posterior end of the metacorpus.

Inserting on the lumen of the isthmus, the narrow stretch of pharynx between the metacorpus and basal bulb, is pm5, a single syncytium with six nuclei. The pm5 syncytium also extends anteriorly in the metacorpus (Fig. 7C), where it has anterior extensions that interdigitate with the pm4 syncytium (Fig. 8C). Posteriorly, pm5 also inserts on the lumen of the anterior basal bulb. Like pm4, pm5 is made up of three radial sectors, each with two adradial nuclei each, although these sectors are joined by bridges between them (Figs 7D, 8E). Each of these bridges occurs between the mc2 cells anteriorly and the mc3 posteriorly.

The other muscles of the basal bulb are pm6, pm7, and pm8. Relative to other pharyngeal muscles, the cells in these classes are poor in myofilaments, which are radially oriented (Fig. 7E). The pm6 and pm7 classes each consist of three, uninucleate radial cells. The pm6 muscles extend as multiple, separated bands to the basal lamina, are each flanked by glands on either side, and insert on the middle of the basal bulb (Fig. 8C, D). The pm7 cells are relatively small and, unlike all other muscle cells of the *P. pacificus* pharynx, as well as pm7 muscles in *C. elegans* and outgroups (Albertson and Thomson, 1976; Zhang and Baldwin, 2000, 2001), do not extend radially to the basal lamina of the pharynx (Fig. 8D, E). The most posterior cell of the pharynx is pm8, a single cell with one nucleus (Figs 7E, 8D, E). This cell forms a cap to the basal bulb and adjoins the pharyngo-intestinal valve posteriorly.

Anchoring the luminal radii of the postcorpus are two sets of non-muscular, marginal cells (Fig. 8A): mc2, which lines the posterior metacorpus, all of the isthmus, and some of the anterior basal bulb (Fig. 7C); mc3, which lines the rest of the basal bulb except for where pm8 forms a cap (Fig. 7D, E). The classes mc2 and mc3 consist of three uninucleate cells each. The left dorsal mc2 nucleus lies in the isthmus, while the other two are in the basal bulb (Fig. 2B). The mc2 cells have tunnels or grooves through which the pm4 syncytial bridges pass (Figs 7C, 8G), although these marginal cells otherwise separate the radial sectors of pm4. Further, the left subdorsal and ventral nuclei are thin and elongated, whereas the right subdorsal nucleus is shorter and round (Fig. 2B). The three mc3 nuclei are symmetrically arranged in the posterior ends of the cells. Like the muscles of the basal bulb, each mc3 cell is flanked on either side by a gland cell. Where each mc2 cell meets an mc3 cell anteriorly along the lumen is an opening for the bridges of the pm5 cell (Fig. 8H).

### 3.5 Putative XXX cell

Because of the embryonic role of XXX cells in *C. elegans* facial development, we searched for possible homologs of these cells in *P. pacificus*. Our search was specifically for isolated cells, without processes, in the head. We identified a single candidate, on the left side of the pharynx (Fig. 2C). No counterpart was observed on the right side, even after making allowance for asymmetry of position. However, data from another individual, Specimen 148 from Bumbarger et al. (2013), suggest that the XXX cells can exist as a bilateral pair (Steven Cook, pers. comm.). The cell we observed is located just posterior to the nerve ring and shows only a short posterior process in the present specimen, although processes are known to be variable for the XXX cells of *C. elegans*. Unlike neurons, which have similarly small cell bodies, the observed cell does not enter the nerve ring. We hypothesize that the free-standing cell is homologous with XXXL of *C. elegans*.

### 3.6 Observations of the *C. elegans* pharynx

The numbers of marginal cells and muscle cells or syncytia of *P. pacificus* were found to be mostly conserved with those of *C. elegans*. Nearly all obvious differences in the cells’ numbers, orientation, or form are correlated with differences in gross morphology. Namely, the two species differ in cells producing anterior feeding structures and the mouth and in cells comprising the basal bulb, which has a grinder in *C. elegans* but lacks one and is mostly glandular in *P. pacificus*. However, two potential differences without obvious correlates in pharyngeal morphology or behavior stood out. We found that pm4 and pm5 of *P. pacificus* are both single syncytia rather than classes of three syncytia each. Because continuous cytoplasm should impact electrical and chemical gradients in muscle function, such a difference might be important for comparative studies of pharyngeal behavior. Yet the cytoplasmic bridges between radial sectors are relatively small, and easy to neglect, parts of those cells. Therefore, we revisited the original TEM images through the metacorpus and isthmus of *C. elegans*, to see whether similar bridges might be detected. Indeed, images of specimens N2T and N2W, which were used for the original TEM reconstructions of the *C. elegans* pharynx, show connections among radial sectors in both pm4 and pm5 (Table S1). As in *P. pacificus*, syncytial connections in pm5 are across the junctions of mc2 and mc3. Unlike *P. pacificus*, in which pm4 connects through holes in the mc2 cells, the sectors of pm4 connect across the junctions of mc1 and mc2 in the metacorpus in *C. elegans*. These observations, guided by our findings in *P. pacificus*, indicate that the metacorpus and isthmus muscles are each a single syncytium in *C. elegans*.

## 4 Discussion

### 4.1 A conserved set of cells produce novel, predatory mouthparts in nematodes

Our reconstruction of the *P. pacificus* feeding apparatus identified all of its components and adjoining cells. As a result, we determined which of those cells have diverged during the evolution of predatory feeding from microbivory. A comparison of *P. pacificus* and *C. elegans* shows that all muscle and epithelial cell classes in the head and pharynx, in terms of both cell fates and number of nuclei, are identical between the two nematodes (Table 1). Rather than requiring any turnover in the number or fates of those cells, divergence between predatory and strictly microbivorous mouthparts was through the cells’ form and their extent of contacts to each other. This divergence specifically occurred in the classes of the pharyngeal cells (especially in pm1 and “e” cells) that line the teeth and inside of the mouth. To infer the direction of change in form, arrangement, and connectivity of cells during the evolution of teeth, we consider states in outgroup (Tylenchina) species (Fig. 9).

**Fig. 9.**
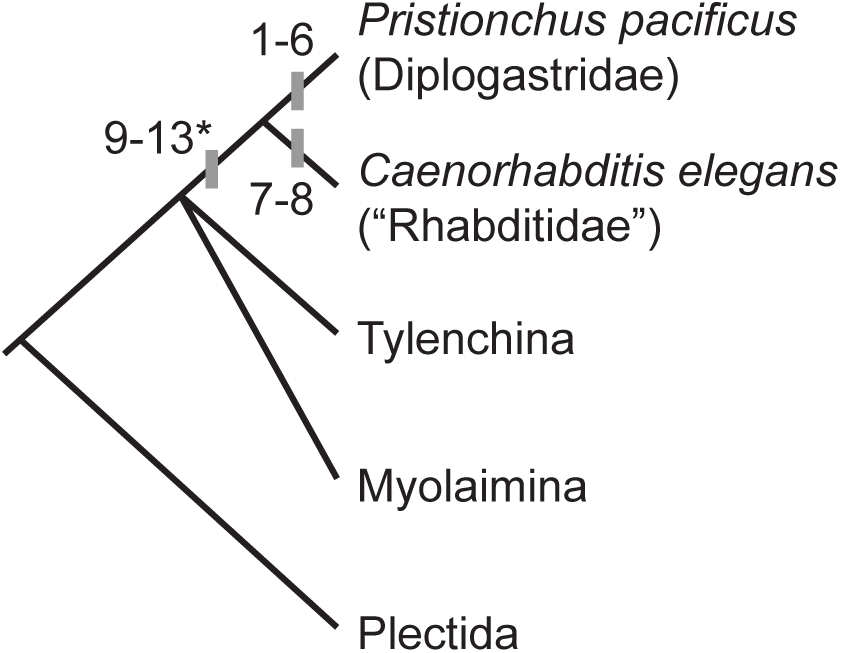
Simplified tree of Rhabditida showing differences in pharynx anatomy within Rhabditina, represented by *Pristionchus pacificus* and *Caenorhabditis elegans*. Character mapping by simple parsimony. Representatives of each taxon listed in Table 1. Exclusive to *P. pacificus* are: (1) left-right asymmetry of pm1; (2) pm1 with longitudinal muscle in radial arms; (3) some pm1 nuclei in procorpus; (4) e1–e3 surrounding and between teeth; (5) telescoped hyp cells; (6) hyp4 producing part of cheilostom. Exclusive to *C. elegans* are: (7) pm2 as three syncytia; (8) mc3 as one syncytium. Shared among Rhabditina representatives, in contrast to Tylenchina and Myolaimina, are: (9) pm1 as three syncytia; (10) pm3 as three syncytia (“paired cells”); (11) pm4 as one syncytium; (12) pm5 as one syncytium; (13) absence of HypD. Asterisk indicates equivocal character polarity, given the unresolved position of Myolaimina (Ahmed & Holovachov, 2021), missing states in Plectida, and (for 13) incomplete data also in Myolaimina.

**Table 1.**
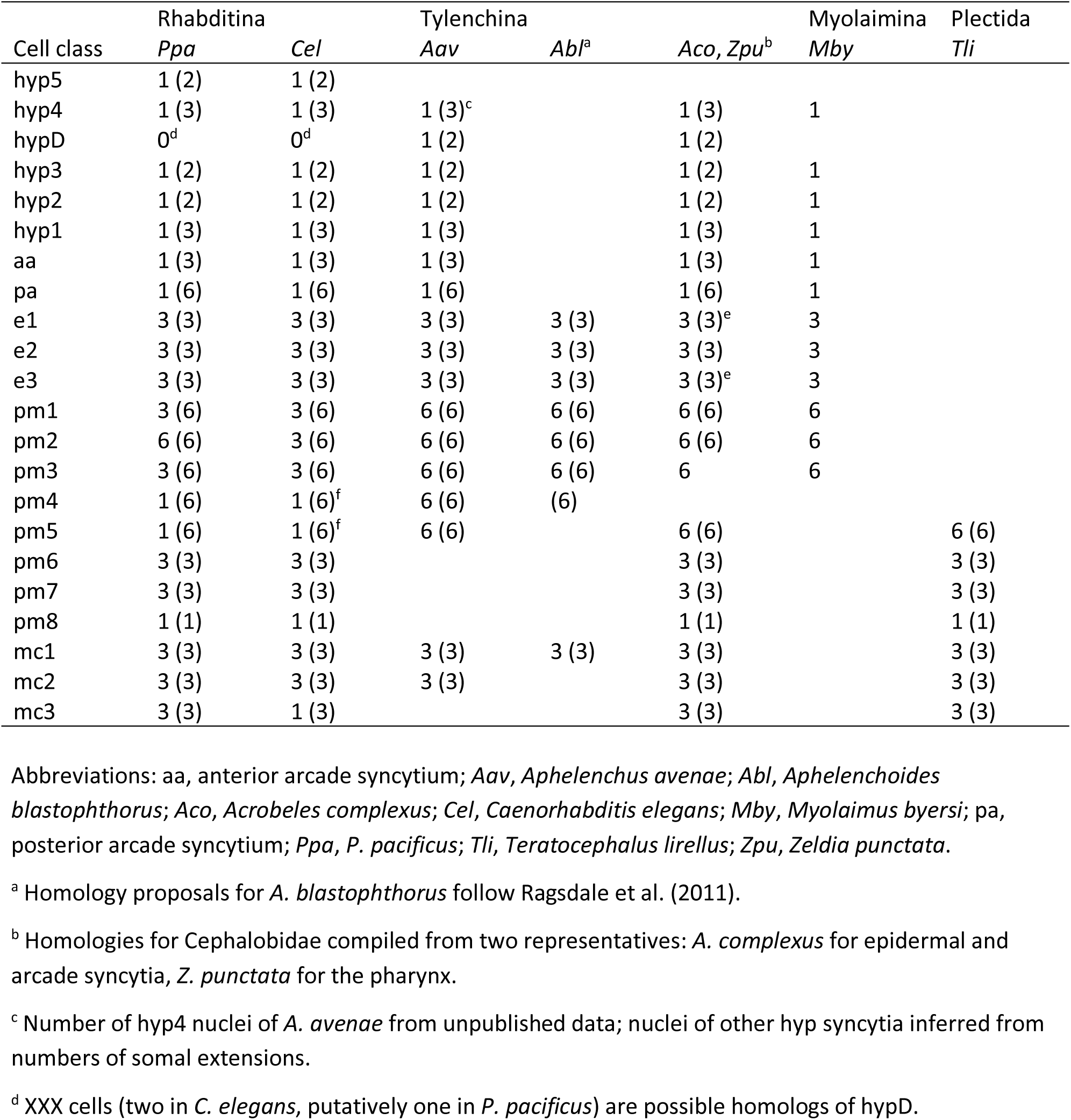
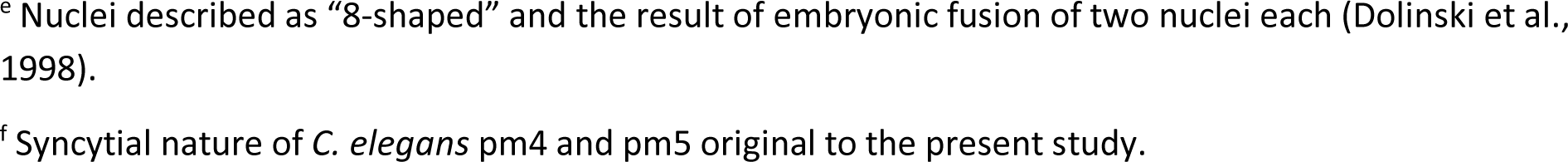
Conservation of numbers of cells/syncytia and, in parentheses, nuclei of putatively homologous epithelial cells of the face and feeding apparatus in *Pristionchus pacificus*, other Rhabditida, and a further outgroup (numbers omitted where unknown or ambiguous).

| Cell class | Rhabditina |  | Tylenchina |  |  | Myolaimina | Plectida |
| --- | --- | --- | --- | --- | --- | --- | --- |
|  | *Ppa* | *Cel* | *Aav* | *Abl*<sup>a</sup> | *Aco, Zpu*<sup>b</sup> | *Mby* | *Tli* |
| hyp5 | 1 (2) | 1 (2) |  |  |  |  |  |
| hyp4 | 1 (3) | 1 (3) | 1 (3)<sup>c</sup> |  | 1 (3) | 1 |  |
| hypD | 0<sup>d</sup> | 0<sup>d</sup> | 1 (2) |  | 1 (2) |  |  |
| hyp3 | 1 (2) | 1 (2) | 1 (2) |  | 1 (2) | 1 |  |
| hyp2 | 1 (2) | 1 (2) | 1 (2) |  | 1 (2) | 1 |  |
| hyp1 | 1 (3) | 1 (3) | 1 (3) |  | 1 (3) | 1 |  |
| aa | 1 (3) | 1 (3) | 1 (3) |  | 1 (3) | 1 |  |
| pa | 1 (6) | 1 (6) | 1 (6) |  | 1 (6) | 1 |  |
| e1 | 3 (3) | 3 (3) | 3 (3) | 3 (3) | 3 (3)<sup>e</sup> | 3 |  |
| e2 | 3 (3) | 3 (3) | 3 (3) | 3 (3) | 3 (3) | 3 |  |
| e3 | 3 (3) | 3 (3) | 3 (3) | 3 (3) | 3 (3)<sup>e</sup> | 3 |  |
| pm1 | 3 (6) | 3 (6) | 6 (6) | 6 (6) | 6 (6) | 6 |  |
| pm2 | 6 (6) | 3 (6) | 6 (6) | 6 (6) | 6 (6) | 6 |  |
| pm3 | 3 (6) | 3 (6) | 6 (6) | 6 (6) | 6 | 6 |  |
| pm4 | 1 (6) | 1 (6)<sup>f</sup> | 6 (6) | (6) |  |  |  |
| pm5 | 1 (6) | 1 (6)<sup>f</sup> | 6 (6) |  | 6 (6) |  | 6 (6) |
| pm6 | 3 (3) | 3 (3) |  |  | 3 (3) |  | 3 (3) |
| pm7 | 3 (3) | 3 (3) |  |  | 3 (3) |  | 3 (3) |
| pm8 | 1 (1) | 1 (1) |  |  | 1 (1) |  | 1 (1) |
| mc1 | 3 (3) | 3 (3) | 3 (3) | 3 (3) | 3 (3) |  | 3 (3) |
| mc2 | 3 (3) | 3 (3) | 3 (3) |  | 3 (3) |  | 3 (3) |
| mc3 | 3 (3) | 1 (3) |  |  | 3 (3) |  | 3 (3) |

Abbreviations: aa, anterior arcade syncytium; *Aav*, *Aphelenchus avenae*; *Abl*, *Aphelenchoides blastophthorus*; *Aco*, *Acrobeles complexus*; *Cel*, *Caenorhabditis elegans*; *Mby*, *Myolaimus byersi*; pa, posterior arcade syncytium; *Ppa*, *P. pacificus*; *Tli*, *Teratocephalus lirellus*; *Zpu*, *Zeldia punctata*.
<sup>a</sup> Homology proposals for *A. blastophthorus* follow Ragsdale et al. (2011).
<sup>b</sup> Homologies for Cephalobidae compiled from two representatives: *A. complexus* for epidermal and arcade syncytia, *Z. punctata* for the pharynx.
<sup>c</sup> Number of hyp4 nuclei of *A. avenae* from unpublished data; nuclei of other hyp syncytia inferred from numbers of somal extensions.
<sup>d</sup> XXX cells (two in *C. elegans*, putatively one in *P. pacificus*) are possible homologs of hypD.
<sup>e</sup> Nuclei described as “8-shaped” and the result of embryonic fusion of two nuclei each (Dolinski et al., 1998).
<sup>f</sup> Syncytial nature of *C. elegans* pm4 and pm5 original to the present study.

The most striking change in form is in pm1, which like *C. elegans* but unlike outgroups consists of three radial syncytia (Burr & Baldwin, 2016). Although the orientation of pharyngeal muscle filaments was previously obvious from light-microscopical observations, two fine-structural features are distinct from both *C. elegans* and Tylenchina outgroups. First, the pm1 syncytia of *P. pacificus* show numerous expansions into neighboring cells (pm3) that pump the procorpus, unlike in either *C. elegans* or the cephalobid *Zeldia punctata* (Dolinski et al., 1998). Second, the form and location of several pm1 nuclei are unique in *P. pacificus*, such that those located in large adradial arms of pm1 muscle are larger and in the procorpus. In contrast, nuclei in thinner cell processes are smaller and located in the metacorpus. This is similar to homologs in *C. elegans* and even in the hypothesized homologs in the stomatostylet-bearing nematode *Aphelenchus avenae* (Ragsdale et al., 2011). The larger arms are those that actuate the teeth, whereas the others insert on immovable structures or, in the case of the right subventral tooth, a part of the tooth oblique to the muscle’s lever. Because the movement of teeth and pharyngeal pumping are coupled (Wilecki et al., 2015; Okumura et al., 2017), we suggest that the arrangement of nuclei and form of the cells reflect function of the novel mouthparts.

Another difference from *C. elegans* and Tylenchina, as well as Myolaimina, is in the epidermis, specifically hyp4. In other species, the hyp4 syncytium lines the anterior body wall cuticle, whereas hyp3, hyp2, and hyp1 form the lips and cheilostom (White, 1988; Bumbarger et al., 2006; Ragsdale et al., 2008; Giblin-Davis et al., 2010). In *P. pacificus*, the hyp4 syncytium likewise lines the anterior body wall, but it also has medial extensions that line the flaps that cover the mouth. Thus, our results challenge the homology of the mouth opening, which in *P. pacificus* corresponds to both the cheilostom and the body-wall cuticle distal to the lips.

Diplogastridae are known for complex morphologies of the “cheilostom,” which includes various forms of rods, plates, and flaps (Fürst von Lieven & Sudhaus, 2000; Kanzaki & Giblin-Davis, 2015). This contrasts with outgroups, in which the lumen of the cheilostom is most often a simple tube. In at least one case, the *triformis* group of *Pristionchus*, cheilostomatal structures even differ within species (Ragsdale et al., 2013a; Herrmann et al., 2019). Based on our findings, we speculate that the origin and divergence of such forms has been mediated by hyp4.

### 4.2 Implications for the origins of moveable teeth

The spatial arrangements of mouth cells also have implications for the evolutionary origins of the teeth themselves. Teeth of various types have arisen multiple times in nematodes, and the moveable teeth of Diplogastridae represent just one of these events (Chitwood & Chitwood, 1950). However, the dorsal tooth in particular does not have clear homology with any one structure in microbivorous nematodes, nor are there any species that have retained transitional forms (Susoy et al., 2015). Nonetheless, the teeth are cuticular structures and likely have origins in some pre-existing cuticle in the mouth. The mouth cuticle of free-living (non-parasitic) Rhabditida can be divided into compartments, or “rhabdia” (Steiner, 1933). Although homologies of some rhabdia among nematode groups were historically debated (*e.g.*, Andrássy, 1962; Goodey, 1963; Poinar, 1971), ultrastructural studies suggest that homologies can be tested robustly by identifying the classes of cells that line the rhabdia (Baldwin & Eddleman, 1995; De Ley et al., 1995). However, some homologies of the diplogastrid mouth were until now unresolved even in light of TEM data, as gathered from *Acrostichus halicti* (Baldwin et al., 1997). Furthermore, the position of the dorsal gland orifice (DGO), which opens into the walls of the mouth between two rhabdia in other nematodes, runs through the dorsal tooth in Diplogastridae, deepening the need to identify rhabdion homologies.

Because the DGO is on the dorsal tooth in Diplogastridae, reconstruction of the pm2 cells can help distinguish between scenarios for the teeth’s origins. Specifically, the presence or absence of pm2 insertions on the teeth would determine if and the extent to which these cells, in addition to pm1, are involved in the secretion of these mouthparts. In all Rhabditida examined so far, the putative homologs of pm2 line a short region of the mouth or pharynx: the telostegostom, which includes the DGO (Albertson & Thomson, 1976; Shepherd et al., 1980; Baldwin & Eddleman, 1995; De Ley et al., 1995; Dolinski & Baldwin, 2003; Giblin-Davis et al., 2010; Ragsdale et al., 2011). In microbivorous species, the telostegostom is the posterior most rhabdion of the mouth. In many species of “Rhabditidae,” including *C. elegans*, the telostegostom is a wall of cuticle posterior to three immoveable flaps (“glottoid apparatus”) that are radially arranged between the apices of the pharyngeal lumen (Sudhaus, 2011). These immoveable flaps are lined by pm1 muscle and are hence part of the metastegostom. Because of this, and based on TEM data from *A. halicti*, it has been hypothesized that the tooth evolved from joining of the dorsal flap and a “bulge” derived from the telostegostom (Fürst von Lieven & Sudhaus, 2000). However, the telostegostom had still remained undefined in Diplogastridae: in *A. halicti*, cells of two classes (“mb,” “mc”) were found to line the tooth, the posterior of which (mc) lined the DGO, while a third class (“md”) consisted of cells lining the area posterior region of the mouth. Thus, diplogastrid mouthparts and associated muscles did not previously have clear homologs in *C. elegans*.

Here, we have resolved that *P. pacificus* has exactly the same number of muscle cell classes and nuclei as *C. elegans* and, therefore, that homologies of the diplogastrid mouth are reconciled with a conserved set of cells. We have found that a single set of muscle cells, pm1, inserts on the teeth. These cells, which have multiple contractile arms per syncytium, correspond to both mb and md in *A. halicti* as pictured in transverse section. Similar to outgroups, but unlike in *C. elegans*, pm2 in *P. pacificus* is a set of three pairs of adradial, uninucleate cells. In the dorsal pair, where myofilaments are sparse, they are oriented to dilate the dorsal gland duct at the DGO. Further, these cells extend posteriad between lobes of the dorsal tooth. Thus, only the pm1 muscles actuate the dorsal tooth itself, where they insert along the entirety of the tooth structure, whereas the pm2 cells extend transversely into the tissue making the rest of the tooth. Given this arrangement, our results support that most, but not necessarily all, of the dorsal tooth arose from the metastegostom. However, whether the cuticle lining the dorsal gland canal connecting the gland duct to the DGO—that is, the cuticle lining the thin ventral margin of the dorsal tooth—is produced by pm2, or whether the pm2 simply redirects the DGO canal to cuticle wholly derived from pm1 is still unresolved. The electron density of the cuticle of the tooth and that of its base, which would in principle distinguish alternative origins of that cuticle, do not obviously favor one hypothesis over the other.

In contrast to the dorsal tooth, in which the gland canal may have involved the co-option of pm2-secreted cuticle into the tooth, the right subventral tooth and left subventral ridge are lined completely by pm1. Instead of joining the tooth, the subventral pm2 cells insert on the walls of the pharyngeal lumen. Together with pm3, which inserts on the lumen in the same transverse plane, pm2 cells likely function to dilate the posterior part of the mouth. Therefore, our reconstruction supports the opposing right subventral tooth, as well as the left subventral ridge, of diplogastrids to be completely of metastegostomatal origin. Consequently, these structures may each have been derived from a subventral flap of a glottoid apparatus as known from non-diplogastrid “Rhabditidae.”

Another difference in spatial arrangement that has apparently enabled teeth to become moveable, a key feature of diplogastrid mouthparts, is in the e1 and e3 cells. In both *C. elegans* and microbivorous outgroups, these cells make up an epithelium that forms a ring of cuticle continuous with the rest of the nematodes’ tube-like mouth (Albertson & Thomson, 1976; Baldwin & Eddleman, 1995; De Ley et al., 1995). Likewise, putative homologs of these cells in microbivorous Tylenchina, which are myoepithelial rather than epithelial, are arranged in rings.

However, in *P. pacificus*, the e1 and e3 cells do not form a simple ring but show longitudinal extensions between pm1 muscles, such that the cells line cuticle that surrounds most of the tooth in each sector. This cuticle is of different biochemical composition than either the tooth or the rest of the pharyngeal lumen. Thus, we hypothesize that these rhabdia (prostegostom and mesostegostom) together comprise flexible hinges allowing the movement of both the dorsal and right subventral teeth.

### 4.3 Cellular architecture of a conspicuously asymmetrical form

The feeding structures of nematodes offer an unusual, if not unique, case study for how symmetry is broken in animals. This is because nematode mouthparts are produced on two superimposed axes of symmetry: the left-right axis, as in most other bilaterians, as well as a radial axis. Outside of the pharynx, most nematode cell classes of more than one cell, most frequently in the nervous system, are organized as bilateral pairs, even those belonging to radial groups, such as the cephalic sensillum neurons and glial cells (White et al., 1986). While this is also the case of the pharyngeal nervous system, the epithelial cells of the pharynx are arranged in truly radial groups of three or six. Fixed left-right asymmetry in radial mouthparts occurs in several nematode groups besides Diplogastridae, including in Odontolaimidae (Order Triplonchida) and in Oncholaimidae and Enchilidiidae (Enoplida), the result of at least two evolutionary events (Lorenzen, 1994; Smythe, 2015). Dorsoventral differences without bilateral asymmetry also occur in the teeth (onchia) of Odontolaimidae, such that dorsoventral differences can be distinguished from bilateral ones. In the other two families, the mouth is often dominated by one subventral onchium, while the other two are similar in size, hinting at a break from radial symmetry *per se*. In *P. pacificus*, clear dorsoventral differences are found, namely in the pm2 cells and in the prominence of the short, central arms of pm1. However, we additionally find departures from strictly radial symmetry in *P. pacificus*. The six radial cells that ultimately form the three pm1 syncytia assume one of only two states in terms of their “cell bodies,” when defined as adradial arm (somal extension) form and nucleus form and position. It is the different combinations of these two cell types, which differ across the three pm1 syncytia, that distinguish both dorsoventral and left-right differences. Thus, fine-structural differences around a radial axis in part define left-right asymmetry in diplogastrid teeth.

Our reconstruction also expands the context for studying molecular causes of asymmetry in nematodes. Several mechanisms for the left-right asymmetry of organ development are known in animals with indeterminate cleavage, such as vertebrates (Levin et al., 1995), and animals with determinate cleavage but not cell constancy, such as flies and snails (Spéder et al., 2006; Kuroda et al., 2009). Some of these mechanisms may also be generalizable and ancestral to bilaterians, such as myosin 1D-directed flow (Tingler et al., 2018). However, others may be peculiar to animals with fixed cell lineages, in that at least part of the mechanism might be cell-autonomous after cell fates have been set. Among nematodes, the molecular bases of asymmetry have been extensively studied in the *C. elegans* nervous system, especially in the sensory neurons ASEL and ASER, which are bilaterally paired cells with asymmetric functions (Hobert, 2014). These two cells, which like most neuron pairs are identical in their later patterns of cell division, originate from different blastomeres, such that they differ in whether they are exposed to an intercellular signal (Notch) early in embryogenesis (Priess, 2005). This signaling transiently activates two transcription factors (TBX-37, TBX-38) in ASEL (Good et al., 2004), which then trigger the activity of a miRNA (*lsy-6*) that ultimately activates an alternative gene regulatory network distinguishing left from right (Cochella & Hobert, 2012).

Is an analogous, early cue also required for left-right asymmetry of the *P. pacificus* procorpus? In *C. elegans*, pharynx development similarly proceeds through the inductions of cells from two different lineages, AB.ara-MS and AB.alp-MS (Hutter & Schnabel, 1994), and these inductions also occur in *P. pacificus* (Vangestel et al., 2008). Because AB.alp contributes to the left pm1 and pm2 cells, which belong to cell classes otherwise descended from AB.ara (Sulston et al., 1983), it is possible that some early signal prior to pharynx induction may be involved in symmetry breaking. Although this and alternative hypotheses must still be speculative, our reconstruction narrows the search for developmental mechanisms for a conspicuous asymmetry of nematode form. Specifically, genetic mutants with symmetrical mouthparts (i.e., no subventral teeth, or paired subventral teeth) or, alternatively, antisymmetrical teeth (i.e., randomly left-right placement of a subventral tooth) could help identify the mechanism whereby, and at what stage of development, pharynx symmetry is broken in this species.

### 4.4 Other comparisons with the feeding apparatus of *C. elegans*

Besides one class of mouthpart-associated cells (pm2) and one class of marginal cells (mc3), all epithelia, including muscles, of the *P. pacificus* pharynx are identical to homologs in *C. elegans* in numbers of syncytia and nuclei. Although pm7 and pm8 were not previously detected in the basal bulb of the diplogastrid *Allodiplogaster* (= *Diplenteron*) sp. (Zhang & Baldwin, 1999), we confirm here that the basal bulb of *P. pacificus* has all muscle cells classes and nuclei present in *C. elegans* as well as representatives of Tylenchina (*Zeldia punctata*, *Basiria gracilis*) and even Plectida (*Teratocephalus lirellus*) (Zhang & Baldwin, 2000, 2001; Baldwin et al., 2001). The pm7 and pm8 muscles of *P. pacificus*, however, are reduced in size compared with *C. elegans*. Instead, the three gland cells occupy a larger share of the basal bulb in both *Allodiplogaster* sp. and *P. pacificus* (Zhang & Baldwin, 1999; Riebesell & Sommer, 2017). An apparent exception to the conservation of syncytium number was that pm4 and pm5, which we found to be one syncytium each, although further inspection of original TEM data for *C. elegans* indicates those cell classes are likewise single syncytia. This contrasts with at least one Tylenchina representative, *A. avenae*, which has six uninucleate pm4 cells (Ragsdale et al., 2011).

The nuclei of mc2, the marginal cells of the isthmus and part of the basal bulb, differ from *C. elegans* by their symmetry. Whereas in *C. elegans* the nuclei are radially symmetrical around the posterior part of the isthmus, in *P. pacificus* each nucleus is unique in its combination of shape and position. In *P. pacificus*, we find that the right subdorsal nucleus is round, whereas the other two nuclei are long and thin; further, the left subdorsal nucleus lies in the isthmus, while the other two are in the basal bulb. Consistent with this finding, the presence of one of the mc2 nuclei in the isthmus can indeed be seen by light microscopy (Bumbarger et al., 2013). However, because our reconstruction was for a single specimen, how variable the left-right asymmetry of either position or shape of the mc2 nuclei is unclear.

Whether this asymmetry is fixed or fluctuating has yet to be determined by increased sampling of these traits.

In terms of cell arrangement, one potential difference from *C. elegans* is the chirality of e2 cells, although we again acknowledge that this may an artefact of reconstructing just one specimen of *P. pacificus*. Like in *C. elegans*, the cell bodies of e2 are offset by 60° from where they line the mouth cuticle. However, the twist is sinistral (clockwise when viewed *en face*) in *C. elegans* and dextral in the *P. pacificus* specimen described herein. Examination of more specimens should distinguish whether this chirality is fixed or variable within the species.

A cell we tentatively identified with reference to *C. elegans*, and which may be of consequence for the *P. pacificus* mouth development, is an XXX cell. In *C. elegans*, the XXX cells (XXXL and XXXR) are of hypodermal origin and, during early embryogenesis, function as hypodermis near the hyp4 and hyp5 cells (Hall & Altun, 2008). These cells later retract to the interior of the body to become free-standing cells of varying position and having an anterior process of varying length (White, 1988). Postembryonically the XXX cells serve a neuroendocrine function, particularly in dauer development (Ohkura et al., 2003; Kimura et al., 2011; Schaedel et al., 2012). Possible homologs of the XXX cells have been proposed in Tylenchina, in which a syncytium of two lateral nuclei (hypD) lies between hyp3 and hyp4 in the adult (Bumbarger et al., 2006; Ragsdale et al., 2009). A reconstruction of neighboring cells in the nerve ring ganglia, particularly from more than one specimen, as well as evidence from gene expression reporters should more accurately identify XXX homologs, as well as their variability, in *P. pacificus*.

### 4.5 An anatomical map for comparative developmental genetics in nematodes

Our reconstruction provides a guide to tracking molecular interactions in and among individual cells in a model for morphological polyphenism. For example, the interpretation of gene reporter constructs in the face, mouth, and pharynx requires knowing both nucleus position and the shapes of cell processes. While some of these cells were previously identified based on presumed conservation with *C. elegans*, others have been more elusive, making it difficult to pinpoint mechanisms at an organismal level. Although we have only reconstructed the feeding apparatus for a eurystomatous (predatory) individual, we expect our anatomical map of cell bodies to describe both the eurystomatous and the stenostomatous morphs. For example, an expression reporter for the polyphenism switch gene *seud-1*/*sult-1* was previously generated for a stenostomatous-biased line of *P. pacificus*, such that both developing juveniles and adults of both morphs could be examined (Bui et al., 2018). All nuclei that can be identified in those specimens, including those of the asymmetric pm1 muscles, are constant between morphs. However, differences in the form of those cells may exist between morphs, and we propose that our reconstruction could help to identify any such differences.

We also expect a resolved anatomy to inform comparative studies of pharyngeal behavior. The *C. elegans* pharynx is well established as a model for the genetics and circuitry of behavior, a benefit of the organ’s relatively simple nervous system (Avery & You, 2012). Consequently, *C. elegans* can serve as an anchor for studies of the evolution of feeding behavior in nematodes (Chiang et al., 2006). Since the connectome of the *P. pacificus* pharynx was described, circuit- and systems-level comparisons between two species with divergent feeding behaviors is possible (Bumbarger et al., 2013). Further, the study of pharyngeal behavior of *P. pacificus* has since been informed by genetic mutants (Ishita et al., 2021). In terms of morphology, similarities and differences in syncytium number between *P. pacificus* and *C. elegans* could help to interpret results of tissue coordination in pharyngeal pumping. They might also inform studies of function from form: for example, two types of nucleated muscle arms are present in the same cell class (pm1) in the same species of nematode, offering a built-in comparison for the structures they affect. Ultimately, the coordination between sensory systems, the central nervous system, and the feeding apparatus should be possible to dissect in this predatory nematode, especially in comparison to the *C. elegans* model. The known anatomy will give an organismal context to those comparisons.

## Acknowledgments

We thank Ralf Sommer for providing the original TEM images used for our reconstruction, and we thank Dan Bumbarger for technical assistance at the start of this project. This work was supported by the US National Science Foundation (IOS-1911688 to E.J.R.).

## Conflict of interests

The authors declare that there are no conflicts of interests.

## Author contributions

E.J.R. conceived and designed the research; C.J.H. and S.M.M. performed the research; C.J.H. and E.J.R. analyzed the data and wrote the manuscript.

## Data availability statement

The data that support the findings of this study have been deposited in Dryad (https://doi.org/10.5061/dryad.z612jm6d4).

## Notes

### Competing Interest Statement

The authors have declared no competing interest.

### Summary of Updates

The manuscript was revised following single-blind peer review.

