## Supplemental Data 1 for "Comparative reconstruction of the predatory feeding structures of the polyphenic nematode *Pristionchus pacificus*"

### Three-dimensional reconstruction of the face and feeding apparatus of the predatory nematode *Pristionchus pacificus*

_____________________

Department of Biology, Indiana University, Bloomington, IN, 47405, USA

**Table S1.** All WormImage (https://www.wormimage.org) image numbers for *C. elegans* N2T, and N2W showing pm4 and pm5 syncytial connections between radial sectors.

| Specimen name | Syncytial bridges shown | Reference number | Image print number |
|---|---|---|---|
| N2T | pm4D-pm4VL | N2T_2619 | 748 |
| N2T | pm4D-pm4VL<br>pm4D-pm4VR | N2T_2623 | 752 |
| N2T | pm4VL-pm4VR<br>pm4D-pm4VR | N2T_2627 | 756 |
| N2T | pm5VL-pm5VR | N2T_1814 | 294 |
| N2T | pm5D-pm5VR<br>pm5VL-pm5VR | N2T_1816 | 296 |
| N2T | pm5D-pm5VL<br>pm5VL-pm5VR | N2T_1820 | 300 |
| N2W | pm4VL-pm4VR | N2W_308539 | 0056 |
| N2W | pm4VL-pm4VR<br>pm4D-pm4VR | N2W_308554 | 0071 |
| N2W | pm4VL-pm4VR<br>pm4D-pm4VR<br>pm4D-pm4VL | N2W_308563 | 0080 |
| N2W | pm5VL-pm5VR<br>pm5D-pm5VR<br>pm5D-pm5VL | N2W_309141 | 658 |
